# Senolytic treatment with fisetin reverses age-related endothelial dysfunction partially mediated by SASP factor CXCL12

**DOI:** 10.1101/2025.08.13.670216

**Authors:** Sophia A. Mahoney, Krystyna Mazan-Mamczarz, Dimitrios Tsitsipatis, Nicholas S. VanDongen, Charnae’ Henry-Smith, Ada N. Okereke, Rachel Munk, Sanna Darvish, Kevin O. Murray, Supriyo De, Myriam Gorospe, Douglas R. Seals, Matthew J. Rossman, Allison B. Herman, Zachary S. Clayton

**Author notes:** **Corresponding Authors:** Zachary S. Clayton, PhD, Assistant Professor, Department of Medicine-Geriatrics, University of Colorado Anschutz Medical Campus, 12631 E 17^th^ Ave., Mail Stop B179, Ste 8111, Aurora, CO 80045, USA, Allison B. Herman, PhD, Stadtman Investigator, Laboratory of Cardiovascular Science, National Institute on Aging Intramural Research Program, National Institutes of Health, 251 Bayview Blvd., Room 09B121, Baltimore, MD 212214, USA. Authors contributed equally (Allison B. Herman, Zachary S. Clayton).

## Abstract

**Background:** Advancing age is the strongest risk factor for cardiovascular diseases (CVDs), primarily due to progressive vascular endothelial dysfunction. Cellular senescence and the senescence-associated secretory phenotype (SASP) contribute to age-related endothelial dysfunction by promoting mitochondrial oxidative stress and inflammation, which reduce nitric oxide (NO) bioavailability. However, the molecular changes in senescent endothelial cells and their role in endothelial dysfunction with aging remain incompletely unclear. As such, in this study we sought to identify the endothelial cell senescence-related signalling pathways, endothelial-derived SASP factors, and their impact on endothelial function with aging.

**Methods:** Single-cell transcriptomics was performed on aortas from young (6 months) and old (27 months) mice with and without *in vivo* senolytic treatment with fisetin (100 mg/kg/day administered in an intermittent dosing paradigm) to characterize endothelial cell senescence and transcript expression changes. Circulating levels of SASP factors were measured to validate transcriptional changes. Plasma exposure and protein addition and inhibiton experiments were conducted in isolated mouse arteries and cultured human endothelial cells to determine the causal role of the circulating SASP milieu and specific SASP factors in mediating endothelial dysfunction and underlying mechanisms-of-action.

**Results:** Senescent endothelial cells exhibited elevated expression of SASP factors, particularly *Cxcl12*, which was reversed by fisetin supplementation, with responses also reflected in circulating CXCL12 concentrations. Plasma from old mice impaired endothelial function by inducing vascular cell senescence, reducing NO, increasing mitochondrial oxidative stress, and promoting endothelial-to-mesenchymal transition—effects partially driven by CXCL12 and prevented by fisetin.

**Conclusions:** These results identify the SASP and CXCL12 as drivers of age-related endothelial dysfunction and establish mechanisms of senolytic intervention with fisetin supplementation.

**NOVELTY AND SIGNFICANCE:** *What is known?:*

- Advancing age is the primary risk factor for cardiovascular disease, in part due to progressive endothelial dysfunction.
- Cellular senescence contributes to age-related endothelial impairment through the secretion of a pro-inflammatory milieu known as the senescence-associated secretory phenotype (SASP), which can affect neighboring cells and tissue function.
- Senolytic compounds selectively eliminate senescent cells and improve vascular function in preclinical models of aging.

*What new information does this article contribute?:*

- Endothelial cells are highly susceptible to senescence with aging *in vivo* and are selectively cleared by senolytic treatment with the natural compound fisetin.
- Single-cell transcriptomic profiling identifies CXCL12 as the most highly upregulated SASP factor in senescent endothelial cells and in circulation with aging, both of which are reversed by senolytic treatment with fisetin.
- We identify a specific circulating SASP factor, CXCL12, as a partial mediator of endothelial dysfunction by inducing mitochondrial oxidative stress, impairing nitric oxide bioavailability, and promoting endothelial-to-mesenchymal transition (Endo, which is restored by senolytic treatment with fisetin.

**Summary:** This study provides novel mechanistic insight into how senescent endothelial cells and their secretory products—particularily CXCL12—contribute to age-related endothelial dysfunction. It further demonstrates that senolytic treatment with fisetin reverses these effects, highlighting a promising translational strategy for targeting vascular aging and preserving endothelial health.

**GRAPHICAL ABSTRACT:** 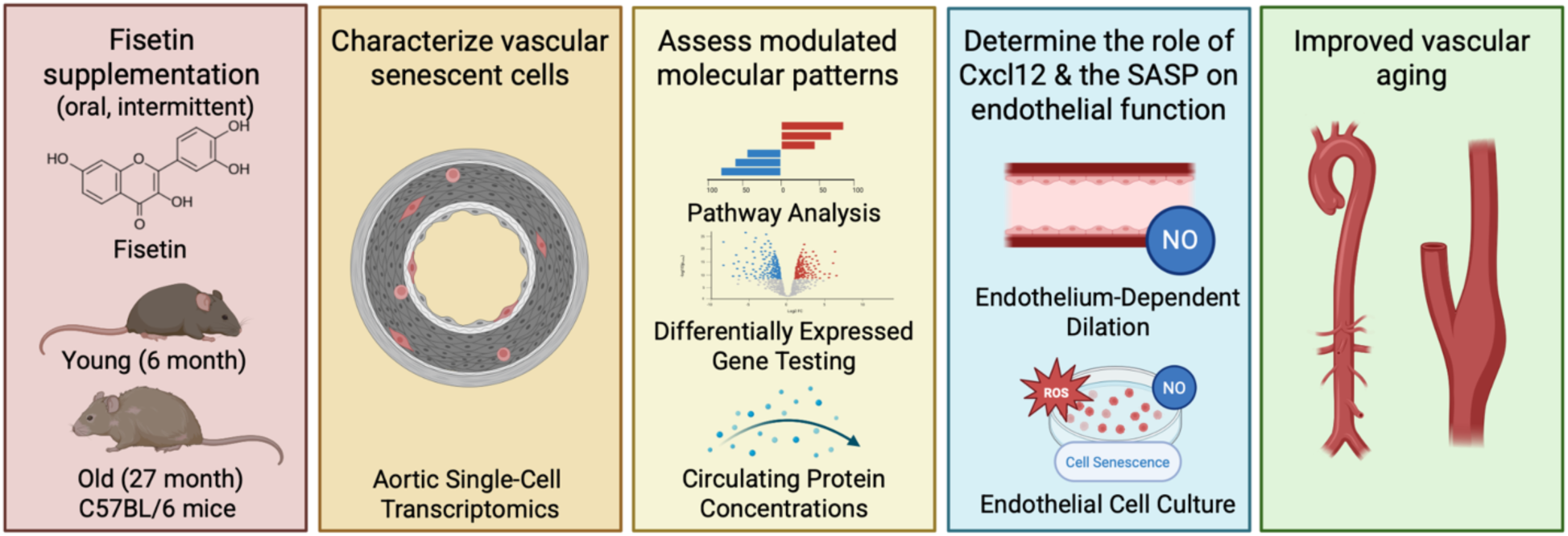

## INTRODUCTION

Advancing age is the primary risk factor for cardiovascular diseases (CVDs), which remain the leading cause of morbidity and mortality worldwide^1,2^. Impaired vascular endothelial function is a key antecedent to the development of clinical CVDs with aging^3^. The antagonistic hallmark of aging, cellular senescence, directly contributes to age-related endothelial dysfunction, in part by promoting chronic inflammation and mitochondrial oxidative stress^4–7^. Despite the adverse effects of cellular senescence on endothelial function, the molecular changes that occur in senescent endothelial cells and the direct mechanisms by which senescent cells induce endothelial dysfunction with advanced age remains unclear.

A putative mechanism by which senescent cells drive physiological dysfunction is through the production and secretion of the circulating senescence-associated secretory phenotype (SASP) milieu, a collection of cytokines, chemokines, proteases, and growth factors^8^. The circulating SASP milieu is a highly heterogenous and dependent on cell type and cellular senescence inducer. Thus, the specific SASP factors that are upregulated with aging in senescent endothelial cells *in vivo* remain to be determined. The circulating SASP milieu may induce cellular and tissue level dysfunction by driving mitochondrial oxidative stress, chronic inflammation, induction of cellular senescence in neighboring cells/tissues, and cell transdifferentiation (i.e., endothelial-to-mesenchymal transition [EndoMT])^8^. Importantly, the circulating SASP milieu is in constant and direct contact with the endothelium, and changes that occur in the circulation contribute directly to age-related endothelial dysfunction^9,10^. As such, endothelial cells may be more susceptible to entering a senescent state given the constant interaction with the circulating SASP milieu^9,11^. However, the circulating SASP factors originating from senescent endothelial cells and their potential impact on endothelial function remains incompletely understood. As such, gaining an endothelial cell-specific understanding of the cellular senescence- and circulating SASP milieu-related molecular mechanisms within the endothelium may be exploited as translational diagnostic biomarkers, and may serve as putative therapeutic targets for novel interventions.

Senolytics – compounds that selectively target and eliminate excess senescent cells^12,13^ – have demonstrated efficacy for suppressing excess vascular cell senescence and improving vascular function in old mice^6,14,15^. Although several senolytic therapies are currently being advanced in preclinical and clinical studies, the natural senolytic fisetin shows efficacy in targeting senescent endothelial cells in culture^16^ and evoking favorable effects on endothelial function with aging^14^, and has an encouraging safety profile for clinical translation^17^. However, the effects of fisetin on senescent endothelial cells and the SASP *in vivo* and how these effects contribute to the benefical effects of fisetin on age-related endothelial dysfunction are incompletely understood.

In the present study, we sought to characterize the vasculature from young (6mo) and old (27mo) mice leveraging *in vivo* fisetin supplementation to explore the molecular, cellular, and physiological mechanisms modulated with aging and senolytic treatment. The central goals of this study were to: 1) assess how aging affects endothelial cell senescence *in vivo* at a single-cell level and identify the key cellular senescence-associated signaling pathways; 2) identify SASP factors upregulated in senescent endothelial cells with aging that may modulate endothelial function; and 3) evaluate the contributions of the circulating SASP milieu and specifc SASP factors in mediating age-related endothelial dysfunction.

We first uncovered that in relation to other vascular cell types, endothelial cells were highly susceptible to becoming senescent with aging and were effectively eliminated by senolytic treatment with fisetin. Within senescent endothelial cells, we identified *Cxcl12* mRNA to be the most highly upregulated with aging and ameliorated with fisetin senolytic treatment, and we observed similar patterns in circulating concentrations of CXCL12 protein. Finally, we revealed a causal role of the circulating SASP milieu in inducing age-related endothelial dysfunction mediated, in part, by the elevated concentration of CXCL12. Overall, these findings distinguish senescent endothelial cells as senolytic targets, establish the circulating SASP milieu as a driver of age-related endothelial dysfunction, and identify CXCL12 as a specific circulating SASP factor that mediates endothelial cell senescence and dysfunction with aging that can be modulated with fisetin senolytic treatment. Further, we uncover mechansims underlying the circulating SASP milieu- and CXCL12-mediated endothelial dysfunction including NO production, mitochondrial oxidative stress, and endothelial cell trandifferentiation.

## METHODS

### Animals

Mouse experiments, including the import, housing, experimental procedures, and euthanasia, were performed strictly under an Animal Study Proposal (ASP #496-LCS-2026) and associated amendments, reviewed and approved by the Animal Care and Use Committee (ACUC) of the National Institute on Aging (NIA). All mice were group-housed at 72°F set point +/-3 under a standard 12 h light/dark cycle and fed ad libitum. Relative humidity was maintained at 30-70%.

For the intervention period, young (6 months) and old (27 months) wildtype mice were randomly assigned to receive vehicle (10% ethanol, 30% PEG400 and 60% Phosal 50 PG) or fisetin (100 mg/kg/day in vehicle). Treatment was administered via oral gavage using an intermittent dosing paradigm – one week on; two weeks off; one week on, as previously established senolytic dosing paradigm **(Figure 1A)**^14,17^.

**Figure 1.**
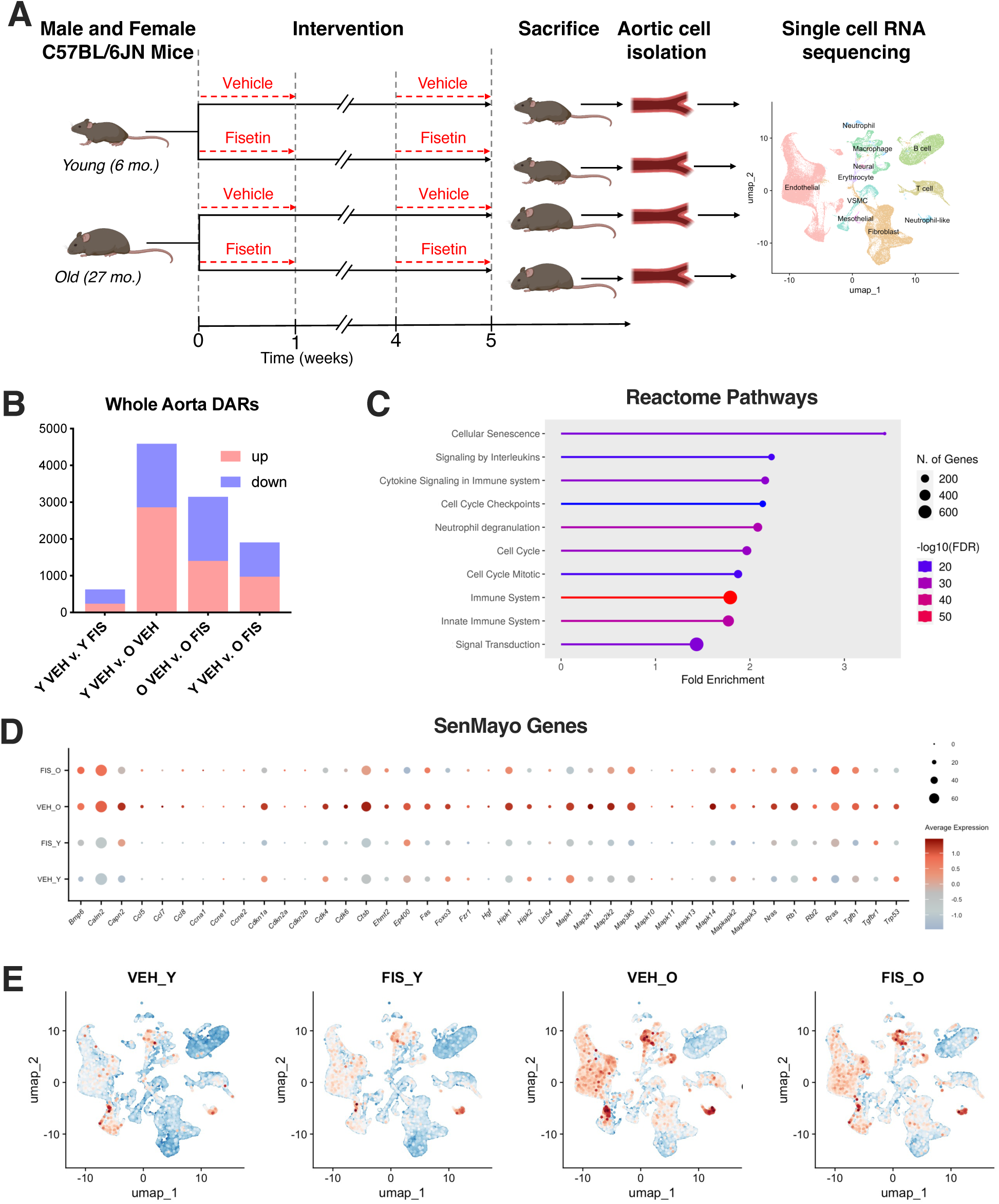
Whole aortic single-cell RNA sequencing identifies senescent cell population with aging that are ameliorated by senolytic treatment with fisetin. Schematic of study design used in animal model resulting in 4 groups: young vehicle (Y VEH), young fisetin (Y FIS), old vehicle (O VEH) and old fisetin (O FIS) treated mice **(A).** Number of diFerentially abundant RNA (DAR) between select treatment groups **(B).** Reactome pathway analysis of increased whole-aorta DARs with age (O VEH v. Y VEH) **(C).** Dot plot analysis of cellular senescence-related DARs **(D).** Uniform manifold approximation and projection (UMAP) of module gene scoring using SenMayo signature to identify senescent cells **(E).** *p<0.05 vs. Y VEH, ^p<0.05 vs. O VEH; triangle represent females, squares represent males.

### Aortic single-cell RNA sequencing

Aortic single-cell transcriptomic analysis was performed, as previously described^18^. Briefly, whole aortas from three mice (per sex and treatment group) were collected, cleaned and digested. Viable cells were sorted using Fluorescence-Activated Cell Sorting for 10x Genomics library preparation. The single-cell libraries were prepared with Chromium Next GEM Single Cell 3’ Kit v3.1 (10x Genomics) with Chromium Next GEM Chip G Single Cell Kit (10x Genomics) according to manufacturer’s protocol with Chromium Controller (10x Genomics). ∼10,000 single cells were used for GEM generation, and the cDNAs were then checked on the Agilent Bioanalyzer with High Sensitivity DNA kit (Agilent). Prepared cDNAs were then used for library preparation that were checked on the Agilent Bioanalyzer with DNA 1000 kit (Agilent Technologies). The libraries were sequenced with Illumina NovaSeq 6000 sequencer at a depth of 70,000-120,000 reads per cell. Sequencing data are deposited in the NCBI’s Gene Expression Omnibus repository GSE296698 (token: yrajmsyujxgjbql).

Single-cell RNA sequencing data were processed using Cell Ranger (version 8.0.1) with the mouse reference reference GRCm39-2024-A (10x Genomics). The obtained read count matrices were subsequently analyzed in R using the Seurat package, version 5.2.0 with default parameters in all functions, unless otherwise specified^19^. Differentially abundant RNA (DAR) testing was performed with the following cutoffs: adjusted p-value < 0.05, average log2 fold change > 0.25, and minimum percentage of cells in either group > 0.1. Pathway analysis was performed using KEGG (version 111.0)^20^ and Reactome (version 90)^21^ and was based on DAR testing.

### Plasma protein quantification

For multiplex analysis, custom murine Luminex Assay kits were designed by R&D Biosystems and executed according to the manufacturer’s instructions

### Plasma-mediated vascular endothelial function

To assess the role of the circulating SASP milieu (plasma) and CXCL12 on endothelial function, an *ex vivo* isolated artery bioasay was leveraged, as previously described^10^. In brief, carotid arteries were excised from young (3-6 month), intervention-naïve wildtype mice and cannulated onto pressure myographs. Plasma collected from young vehicle, old vehicle and old fisetin treated mice was diluted to 5% and perfused luminally through the pressurized arteries for 24 h prior to assessing endothelial function. Following plasma perfusion, endothelial function was measured endothelium-dependent dilation (EDD) and endothelium-independent dilation (EID) in response to increasing doses of acetylcholine (ACh) and sodium nitroprusside (SNP), respectively, as described previously^10,14^. All dose-response data are presented as percent dilation relative to maximum diameter to account for differences in baseline vessel diameter.

### Cultured endothelial cells

Human aortic endothelial cells (HAECs) were grown in basal media supplemented with 5% plasma collected from young, old vehicle, and old fisetin-treated mice. Endothelial cell nitric oxide (NO) and mitochondrial superoxide bioactivity were assessed after 2 h plasma incubation by staining with the fluorescent probe Hoechst (nuclei stain; Thermo Fisher) and either 10 µM diaminorhodamine-4M AM (DAR-4M AM; Sigma-Aldrich; to quantify NO production) for 45 min or 5 µM MitoSOX (Thermo Fisher; to quantify mitochondrial superoxide bioactivity) for 30 min. Senescence-associated β-galactosidase (SA-β-Gal) staining and RNA extractions for gene expression were performed after 24 h of plasma exposure.

### CXCL12 protein addition and inhibiton

Recombinant mouse CXCL12 protein (R&D Systems, Cat. #460-SD-050/CF) in sterile PBS was added back to plasma from young and old fisetin-treated mice, such that the concentration of CXCL12 matched the average levels in old vehicle-treated mice (3210 pg/ml) for CXCL12 add-back experiments. 1 µg/ml of LIT-927 (Selleckchem, Cat. #S8813)^22^ in DMSO was added to the old vehicle plasma for CXCL12 inhibtion experiments.

## RESULTS

### Whole aortic single-cell transcriptomics identifies senescent cell population with aging that are ameliorated with senolytic treatment

To gain insights into the vascular transcriptional changes that occur with aging, we leveraged single-cell RNA sequencing (RNAseq) to compare aortas from young (6mo) and old (27mo) C57BL/6N mice. To study the effects of *in vivo* senolytic treatment on the vasculature, we supplemented young and old mice with vehicle or fisetin, as previously described^14^ **(Figure 1A)**. Single-cell RNAseq workflows allow for the characterization of transcriptomic profiles from distinct cell types of the aorta to identify cell type-specific changes with aging^23^. Unsupervised clustering of the merged samples revealed 11 distinct cell types in the aorta **(Figure 1A** and **Figure S1)**.

We next performed whole-aorta pseudo-bulk RNAseq to determine the general transcriptomic changes with aging and senolytic treatment using differentially abundant RNA (DAR) analysis. Age-related differences (young v. old vehicle) revealed 4589 DARs (2859 higher, 1730 lower)**(Figure 1B)**. Fisetin supplementation in old mice unveiled 3143 DARs with more reduced transcripts than elevated ones (1401 higher, 1742 lower) compared to old vehicle **(Figure 1B)**. Old fisetin mice had a more similar transcriptomic profile to young vehicle mice (974 higher, 931 lower) compared to old vehicle mice **(Figure 1B)**. The young groups (young vehicle v. young fisetin) were the least different with 628 DARs (240 higher, 388 lower)**(Figure 1B)**. Reactome pathway analysis revealed cellular senescence as the most highly upregulated pathway with aging, followed by pathways related to immune dysregulation and cell cycle arrest **(Figure 1C)**. To gain insight into the cell senescence-related transcripts in the vasculature that were modulated with aging and senolytic treatment, we assessed genes in the holistic senescent cell signature SenMayo^24^ that were upregulated with aging (old vehicle v. young vehicle) and downregulated with senolytic treatment (old fisetin v. old vehicle). We found differences in transcripts involved in the SASP, cell cycle arrest maintenance, and proinflammatory signaling **(Figure 1D).**

Given the changes in cellular senescence pathways observed in the vasculature with age, we next sought to assess how endothleial cells were affected relative to other vascular cell types. Leveraging module scoring of SenMayo, we identified senescent cell populations in several vascular cell types including endothelial, macrophage, and neutrophil-like cell populations in aortas from old vehicle mice **(Figure 1E)**. In the aortas from old mice, fisetin had the greatest effects on the SenMayo signature in the endothelial cell population, reinforcing the senolytic effects of fisetin on senescent endothelial cell burden with aging **(Figure 1E)**. Marginal differences were observed on the SenMayo signature in the young groups suggesting minimal effects of fisetin on young, healthy mice **(Figure 1E)**. Together, these data demonstrate a transcriptional shift in the vasculature with aging and suggests that senolytic treatment with fisetin in old mice returns the transcriptome towards a young aortic cell composition by mitigating vascular cell senescence burden, particularly in the endothelium.

### Senescent endothelial cell burden increases in the aorta with aging and is ameliorated by senolytic treatment

We next performed a targeted interrogation of the cellular senescence-related changes in the vascular endothelial cell population with aging and senolytic treatment. To isolate endothelial cell-specific changes, we performed unsupervised clustering of the endothelial cell cluster which produced 12 unique subclusters based on shared transcript expression patterns **(Figure 2A)**. Endothelial cell composition revealed minimal differences between treatment groups in the majority of subclusters, with the exception of subcluster 10 that increased with age (10.7-fold, p<0.0001) and reduced (0.02-fold, p<0.0001) back to young levels, in old mice fisetin **(Figure 2B)**.

**Figure 2.**
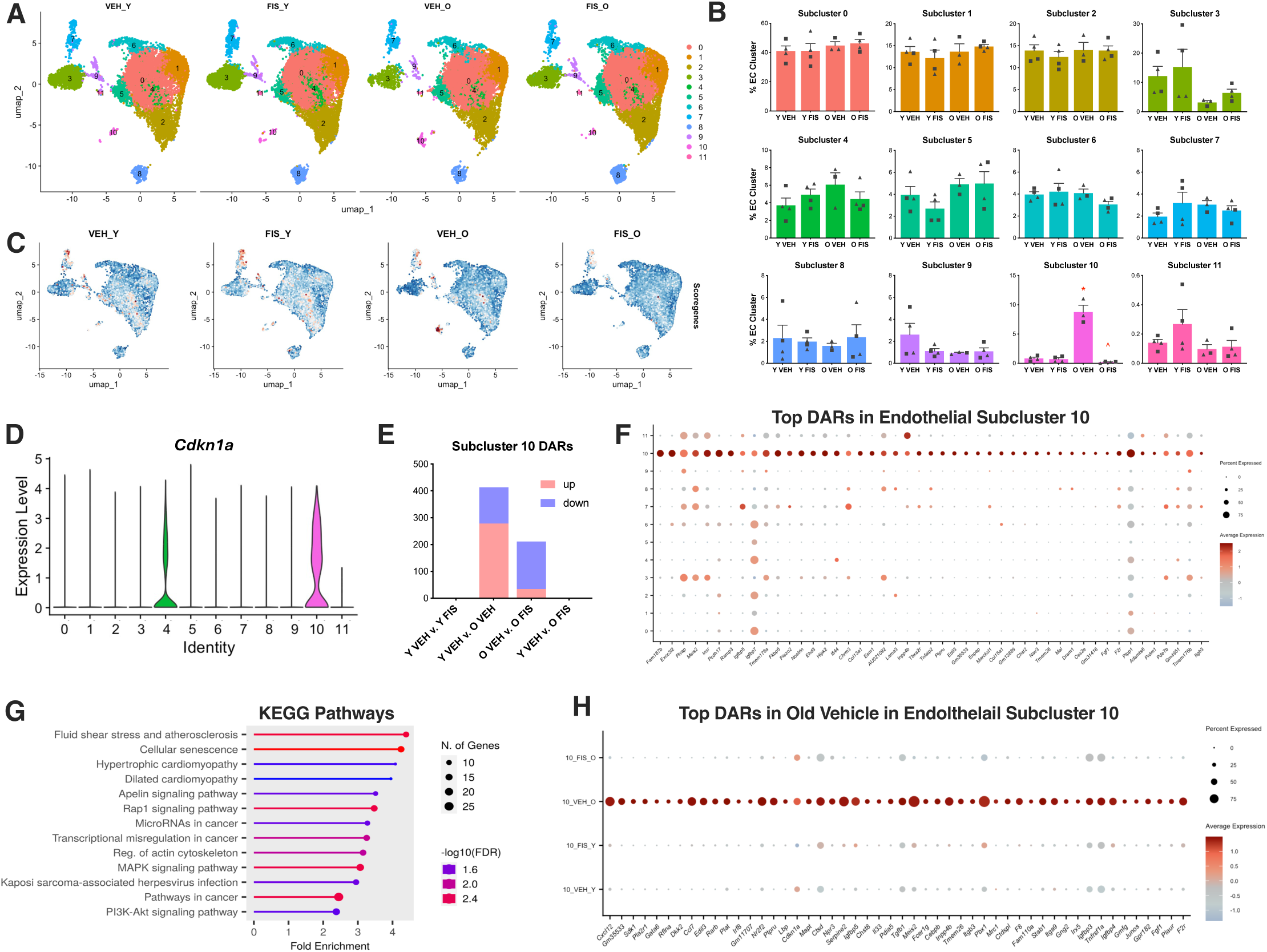
Senescent endothelial cells increase in aortas with aging and are ameliorated by senolytic treatment with fisetin. Uniform manifold approximation and projection (UMAP) **(A)** and subcluster percentage **(B)** of aortic endothelial cells (EC) from the young vehicle (Y VEH), young fisetin (Y FIS), old vehicle (O VEH) and old fisetin (O FIS) treated mice. Module gene scoring using EndoSen signature to identify senescent endothelial cells **(C)**. Violin plot of the canonical cellular senescence marker *Cdkn1a* across clusters on endothelial cells **(D)**. DiFerentially abundant RNA (DAR) between select treatment groups in senescent endothelial subcluster 10 **(E)**. Dot plot analysis of top 50 diFerentially abundant RNAs (DARs) in endothelial subcluster 10 v. other endothelial subcluster **(F)**. KEGG pathway analysis of increased DARs in senescent endothelial subcluster 10 v. other endothelial subcluster **(G)**. Dot plot analysis of the top 50 DARs in endothelial subcluster 10 in old vehicle v. other treatment groups **(H)**. *p<0.05 vs. Y VEH, ^p<0.05 vs. O VEH; triangle represent females, squares represent males.

We next leveraged the endothelial cell-specific senescent signature EndoSen^25^ to determine which subclusters were enriched by cellular senescence-associated signalling pathways. EndoSen module scoring revealed cellular senescence enrichment in endothelial subclusters 4, 7, 9, and 10 with the highest expression being in subcluster 10 **(Figure 2C)**. Moreover, endothelial subcluster 10 had the highest mRNA expression of the canonical cellular senescence marker *Cdkn1a* compared to the other subclusters **(Figure 2D)**.

To further characterize the senescent endothelial cells, we leveraged DAR testing on subcluster 10 to identify gene targets that are modulated with aging and senolytic treatment. Age-related differences (old v. young vehicle) revealed 435 DARs (278 increased, 135 reduced), suggesting a large impact of aging on senescent endothelial subcluster 10 **(Figure 2E)**. Fisetin supplementation in old mice unveiled 211 DARs with more reduced than elevated transcripts (34 increased, 177 decreased)**(Figure 2E)**. Notably, no transcripts were significantly modulated between old fisetin and young vehicle or between the young mouse groups, suggesting that senescent endothelial cells from old mice that received senolytic treatment have a similar transcriptomic profile to that of endothelial cells from young mice and that fisetin has minimal effects on young, healthy endothelial cells **(Figure 2E)**.

The top increased DARs that distinguished the putative senescent subcluster 10 from other endothelial subclusters represented transcripts involved in extracellular matrix remodeling, stress signaling, cell cycle maintenance, and the SASP **(Figure 2F)**. Pathway analysis of the DARs from senescent endothelial subcluster 10 revealed pathways involved in cardiovascular pathology (*i.e.,* atherosclerosis and cardiomyopathy), cellular senescence, cancer, and a general dysregulation of signaling pathways **(Figure 2G)**. Finally, we next identified the top 50 DARs that distinguished the old vehicle group from the other treatment conditions, representing transcripts that were most increased with aging and lower with fisetin supplementation in endothelial subcluster 10 **(Figure 2H)**. We found that these transcripts largely encompass canonical senescence markers (e.g., *Cdkn1a*) and SASP factors. Together, these findings demonstrate increased expression of cellular senescence- and SASP-related signaling pathways in endothelial cells with aging that were ameliorated with senolytic treatment with fisetin.

### Senolytic treatment mitigates the endothelial-derived and circulating SASP in old mice

The SASP represents a collection of pro-inflammatory molecules including cytokines, chemokines, proteases, and growth factors that have varied roles in modulating physiological function^8,26^. Our next aim was to identify candidate circulating SASP factors based on those differentially expressed in the endothelium, particularly those driving the patterns observed in senescent endothelial cell subcluster 10. In subcluster 10, we observed several SASP factors with increased transcripts with age (young vehicle v. old vehicle) that were reduced with fisetin (old vehicle v. old fisetin) including *Ccl4, Ccl7*, *Timp4*, *Igfbp3*, *S100a9*, *Ccl8*, *Fgf1*, *Gdf15*, *Serpine1*, and *Cxcl12* mRNAs **(Figure 3A).**

**Figure 3.**
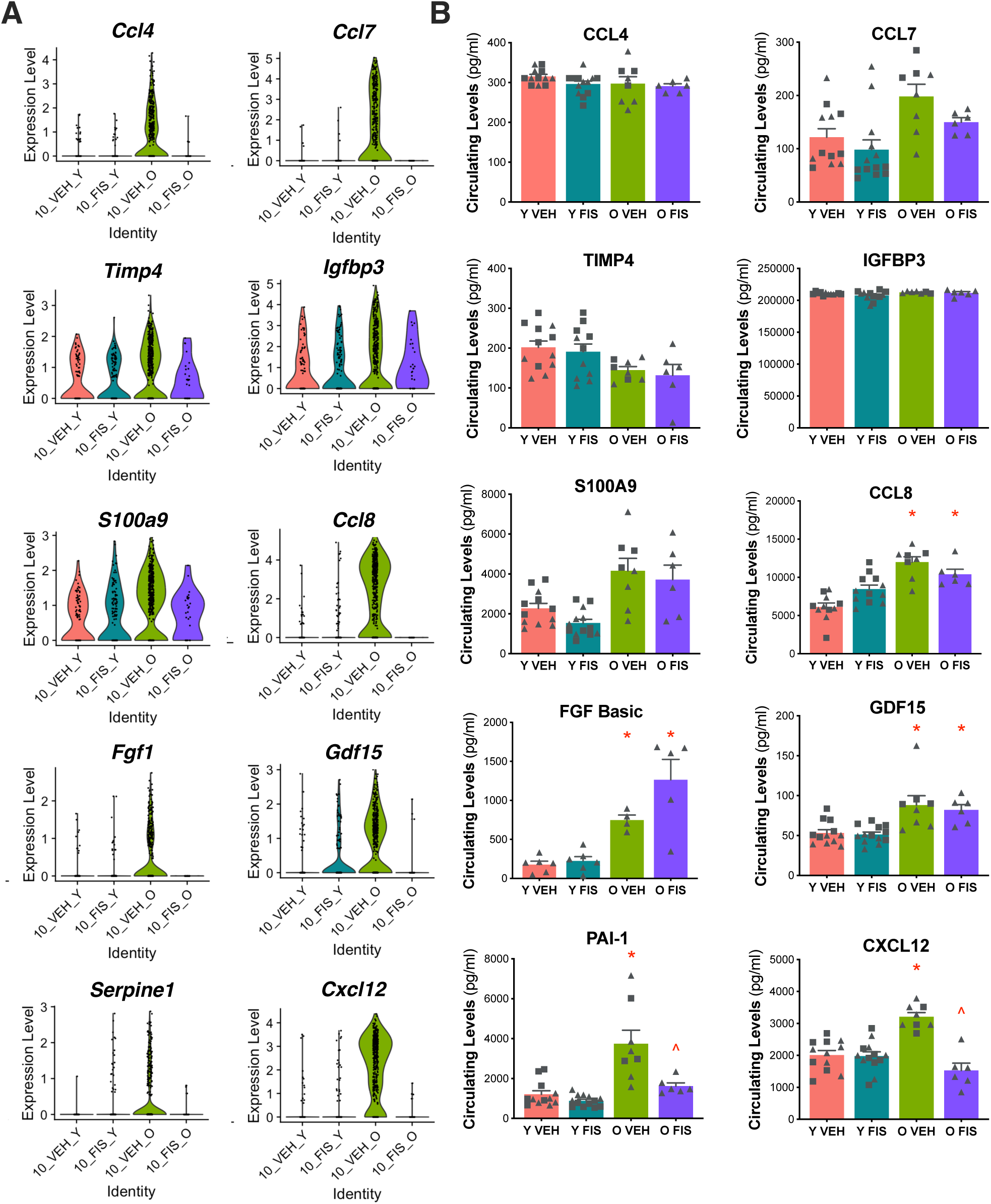
Endothelial and circulating senescence-associated secretory phenotype (SASP) factors increase with age and are ameliorated by senolytic treatment with fisetin. Violin plots representing the top SASP-related diFerentially abundant RNAs (DARs) in senescent endothelial subcluster 10 from the young vehicle (Y VEH), young fisetin (Y FIS), old vehicle (O VEH) and old fisetin (O FIS) treated mice **(A)**. Circulating protein abundance of top SASP-related DARs in plasma **(B)**. *p<0.0001 vs. Y VEH, ^p<0.0001 vs. O VEH; triangle represent females, squares represent males.

Provided the changes observed in mRNAs encoding SASP factors from senescent endothelial cells, we sought to determine if these factors were translated and secreted into the circulating plasma. We selected SASP proteins to measure based on: *a)* the magnitude of age-related differences in senescent endothelial transcript expression and whether these differences with aging were influenced by fisetin supplementation; and *b)* for biological plausibility of altering endothelial function. We measured the protein concentration of candidate SASP factors in the plasma from treated animals. Among these factors, higher concentrations with aging were observed in CCL8, FGF Basic, S100a9, PAI-1, and CXCL12 when comparing young vehicle- and old vehicle mice **(Figure 3B)**. Notably, the elevated PAI-1 and CXCL12 protein expression with aging was ameliorated by fisetin supplementation (old fisetin v. old vehicle)**(Figure 3B)**.

PAI-1, a platelet protein that inhibits fibrin degradation^27^, is a well-established marker of cellular senescence^28^. CXCL12 is a chemokine that regulates angiogenesis^29^ and is implicated in atherosclerotic progression^30^. Given the novelty and putative biological plausibility of CXCL12 in regulating endothelial function (whereas PAI-1 primarily effects platelets), we next sought to investigate whether CXCL12 concentrations were directly implicated in age-related endothelial dysfunction transduced by the circulating SASP milieu and if mitigation of elevated CXCL12 was a mechanism by which fisetin supplementation elicited its benefits on endothelial function with aging.

### Senolytic treatment attenuates circulating SASP milieu-induced endothelial dysfunction with aging

#### Circulating SASP milieu- and CXCL12-induced endothelial dysfunction

Given the observation that select circulating SASP factors changed with aging and senolytic treatment with fisetin, we next aimed to evaluate the functional role of the circulating SASP environment on age-related endothelial dysfunction and the specific effects of CXCL12 as a candidate circulating SASP factor that might be responsible for transducing the effect of the circulating environment on endothelial function. We first tested the effects of the plasma from the young vehicle, old vehicle, and old fisetin groups on endothelial function using carotid artery endothelium-dependent dilation (EDD) following intraluminal plasma perfusion. Average peak EDD was lower following exposure to plasma from old vehicle mice compared to young mice (-17%; p<0.0001), suggesting that plasma from old mice directly induces endothelial dysfunction **(Figure 4A** and **Figure S2B)**. Peak EDD was 14% higher following exposure to plasma from old fisetin mice compared to plasma from old vehicle mice (p=0.001)**(Figure 4A** and **Figure S2A** and **S2B)**.

**Figure 4.**
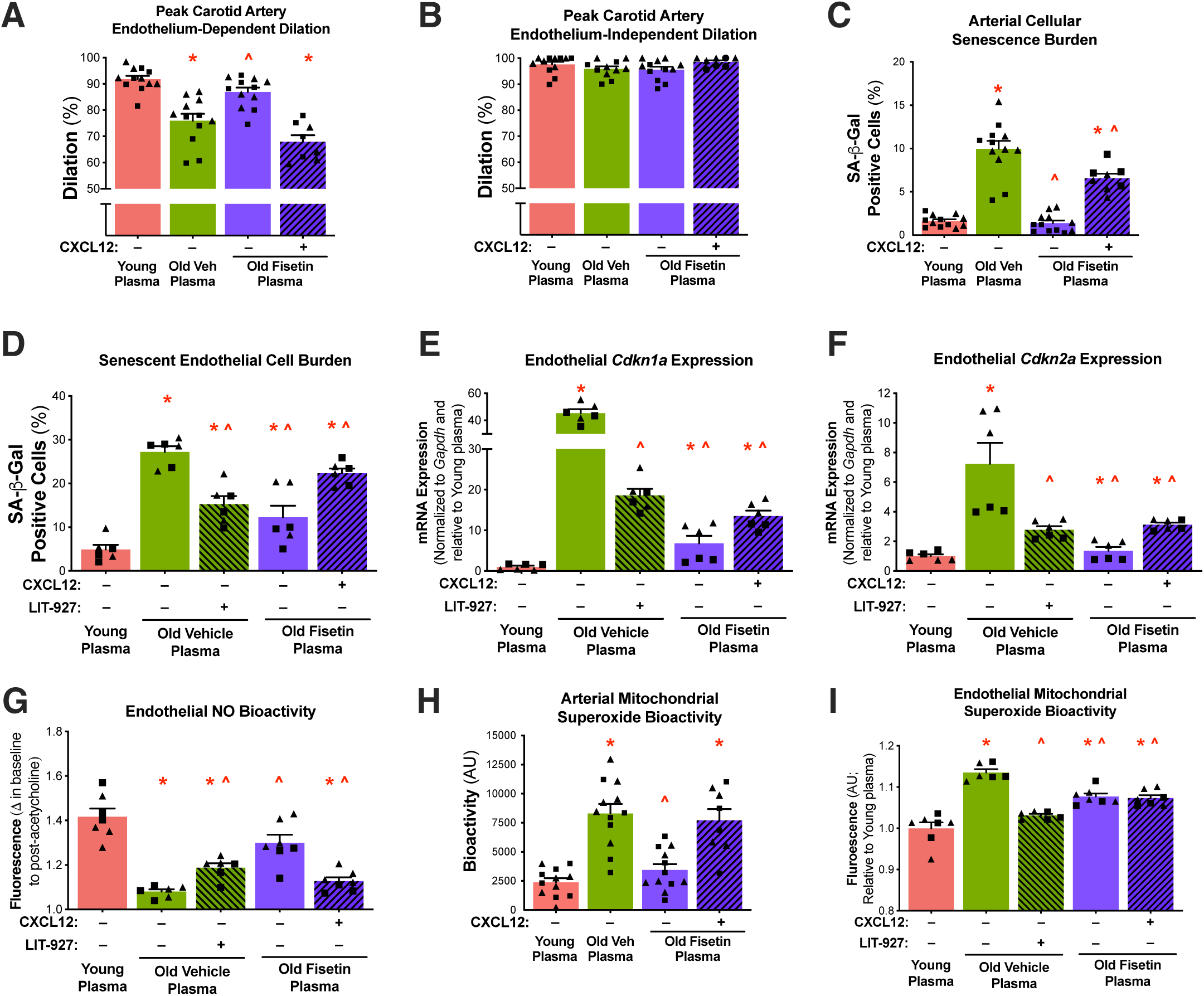
The circulating SASP- and CXCL12-induced endothelial dysfunction with aging and prevention by senolytic treatment with fisetin and CXCL12 inhibition. Endothelium-dependent dilation **(A)** and endothelium-independent dilation **(B)** in isolated carotid arteries from young mice following plasma perfusion with and without the addition of recombinant mouse Cxcl12. Senescence-associated β-Galactosidase (SA-β-Gal) signal following plasma exposure with and without the addition of recombinant CXCL12 and its inhibitor, LIT-927 in isolated arteries **(C)** and human aortic endothelial cells (HAECs) **(D)**. Cellular senescence biomarkers *Cdkn1a* (E) and *Cdkn2a* (F) mRNA in HAECs following plasma exposure with and without the addition of recombinant CXCL12 and LIT-927. Nitric oxide bioactivity in human aortic endothelial cells (HAECs) following plasma exposure with and without the addition of recombinant CXCL12 and LIT-927 **(G)**. Mitochondrial superoxide bioactivity following plasma exposure with and without the addition of recombinant CXCL12 and LIT-927 in isolated arteries **(H)** and HAECs **(I).** Values represent mean ± SEM; *p<0.05 vs. young control, ^p<0.05 vs. old vehicle; triangles represent females, squares represent males.

To determine the mechanistic role of CXCL12 in circulating SASP milieu-mediated endothelial dysfunction, we added back recombinant mouse CXCL12 to old fisetin plasma such that the average concentrations of plasma CXCL12 was similar to that of old vehicle plasma. The addition of CXCL12 to old fisetin plasma reduced peak EDD compared to old fisetin plasma alone (-22%, p<0.0001), ameliorating differences with the old vehicle group (p=0.465)**(Figure 4A** and **Figure S2A** and **S2B)**. No differences in endothelium-independent dilation were observed between any of the groups or conditions (p=0.342), indicating that differences observed in response to plasma exposure occurred in an endothelium-dependent manner **(Figure 4B** and **Figure S2C** and **S2D)**. Together, these data suggest that the circulating SASP milieu underlies age-related endothelial dysfunction and its detrimental effects are in part mediated by CXCL12 and largely reversed by senolytic treatment with fisetin.

#### Circulating SASP- and CXCL12-induced vascular cell senescence

One mechanism by which the SASP elicits its adverse effects is through a humoral-based induction of cellular senescence^6,26^. As such, to investigate the mechanisms by which the circulating SASP modulates endothelial function, we sought to assess SASP-induced cellular senescence by measuring the canonical marker of senescent cell burden senescence-associated β-galactosidase (SA-β-Gal) intensity^31,32^ in arteries (aortic rings) exposed to plasma from young vehicle, old vehicle, and old fisetin mice. We found that when normalized to total cell count there was an increase in SA-β-Gal positive cells in arteries exposed to old versus young plasma (6.1-fold, p<0.0001)**(Figure 4C** and **Figure S3A)**. SA-β-Gal positive cells were lower in arteries exposed to plasma from old fisetin-treated mice compared to old vehicle mice (-86%, p<0.0001)**(Figure 4C** and **Figure S3A)**. The addition of exogenous CXCL12 to old fisetin plasma plasma increased arterial SA-β-Gal positive cells (4.8-fold, p<0.0001) compared to old fisetin plasma alone **(Figure 4D** and **Figure S3A)**.

To isolate the effects of plasma exposure on endothelial cells, we exposed plasma from the treated animals to cultured human aortic endothelial cells (HAECs). Similar to arteries, HAECs exposed to old vehicle plasma had greater SA-β-Gal intensity than HAECs exposed to young vehicle plasma (5.6-fold, p<0.0001)**(Figure 4D** and **Figure S3B)**. SA-β-Gal positive cells were lower in HAECs exposed to plasma from old fisetin-treated mice compared to old vehicle mice (-56%, p<0.0001)**(Figure 4D** and **Figure S3B)**. Inhibition of CXCL12 with LIT-927 (a neutraligand of CXCL12^33^) in old vehicle plasma reduced SA-β-Gal positive cells (-44%, p<0.0001) compared to old vehicle plasma alone **(Figure 4D** and **Figure S3B)**. The addition of exogenous CXCL12 to plasma increased SA-β-Gal positive cells in HAECs exposed to old fisetin plasma (82%, p<0.0001) compared to old fisetin plasma alone **(Figure 4D** and **Figure S3B).**

In addition to SA-β-Gal activity, we confirmed the effects of the SASP and CXCL12 on cellular senescence through the endothelial mRNA expression of the canonical cellular senescence biomarkers *Cdkn1a* and *Cdkn2a.* We found that HAECs exposed to plasma from old vehicle mice had higher *Cdkn1a* (45-fold, p<0.0001) and *Cdkn2a* (6.2-fold, p<0.0001) mRNA expression compared to young plasma **(Figure 4E** and **4F** and **Figure S3A)**. Relative to old vehicle plasma exposure, old fisetin plasma exposure resulted in lower *Cdkn1a* (-85%, p<0.0001) and *Cdkn2a* (-57%, p<0.0001) mRNA expression **(Figure 4E** and **4F** and **Figure S3C** and **S3D)**. The addition of LIT-927 to old vehicle plasma reduced *Cdkn1a* (-59%, p<0.0001) and *Cdkn2a* (-62%, p<0.0001) mRNA expression compared to old vehicle plasma alone **(Figure 4E** and **4F** and **Figure S3C** and **S3D)**. The addition of CXCL12 to old fisetin plasma increased *Cdkn1a* (100%, p<0.0001) and *Cdkn2a* (128%, p<0.0001) mRNA expression compared to old fisetin plasma alone **(Figure 4E** and **4F** and **Figure S3C** and **S3D)**. In combination, these data indicate that the circulating SASP milieu directly induces vascular endothelial cell senescence, in part through CXCL12, and that circulating SASP milieu-induced senescence with aging may be prevented by fisetin supplementation.

#### Circulating SASP- and CXCL12-induced NO production and mitochondrial superoxide bioactivity

Age-related endothelial dysfunction is primarily mediated by reduced nitric oxide (NO) bioavailability, which is a key feature of senescent endothelial cells^34^. Therefore, we assessed NO production in cultured endothelial cells exposed to plasma, examining the effects of both the addition and inhibition of CXCL12. This allowed us to determine the direct role of CXCL12 in mediating differences in endothelial cell NO production induced by the circulating SASP milieu in the context of aging and fisetin supplementation. HAECs exposed to old vehicle plasma exhibited lower NO production compared to young plasma (-24%, p<0.0001)**(Figure 4G** and **Figure S4A)**. This age-related impairment in NO production was prevented in HAECs exposed to old fisetin plasma (20% v. old vehicle plasma, p<0.0001)**(Figure 4G** and **Figure S4A)**. Compared to plasma alone, the addition of exogenous CXCL12 impaired NO production in HAECs exposed to young vehicle (-20%, p<0.0001) and old fisetin (-13%, p<0.0001) plasma **(Figure 4G** and **Figure S4A)**. Further, CXCL12 inhibition by LIT-927 increased NO production in HAECs exposed to old vehicle plasma (10%, p<0.0001)**(Figure 4G** and **Figure S4A)**.

Reduced NO production may occur as a result of mitochondrial oxidative stress, as excessive superoxide scavenges NO to reduce its bioavailability^35^. Thus, we sought to determine whether the circulating SASP milieu increased mitochondrial superoxide bioactivity and if this was mediated by CXCL12. To do so, we next measured mitochondrial-derived superoxide bioactivity in isolated arteries **(Figure 4H** and **Figure S4B)** and cultured HAECs **(Figure 4I** and **Figure S4C)** following plasma exposure with the addition and inhibition of CXCL12. Arterial mitochondrial superoxide bioactivity was higher in aortas exposed to old vehicle plasma compared to young plasma (3.5-fold, p<0.0001)**(Figure 4H** and **Figure S4B)**. Mitochondrial superoxide bioactivity was lower in arteries exposed to plasma from old fisetin-treated mice compared to old vehicle-treated mice (-58%, p<0.0001)**(Figure 4H** and **Figure S4B)**. Compared to old fisetin plasma alone, the addition of exogenous CXCL12 increased mitochondrial superoxide bioactivity (2.2-fold, p=0.001)**(Figure 4H** and **Figure S4B)**.

Similar to arteries, HAECs exposed to old vehicle plasma had greater basal mitochondrial superoxide bioactivity than young plasma (14%, p<0.0001)**(Figure 4I** and **Figure S4C)**. Mitochondrial superoxide bioactivity was lower in HAECs exposed to plasma from old fisetin-treated mice compared to old vehicle mice (-6%, p<0.0001)**(Figure 4I** and **Figure S4C)**. CXCL12 inhibition with LIT-927 reduced mitochondrial superoxide bioactivity in HAECs exposed to old vehicle plasma (-10%, p<0.0001)**(Figure 4I** and **Figure S4C)**. Exogenous CXCL12 induced mitochondrial superoxide bioactivity in HAECs exposed to old fisetin plasma (5%, p=0.029)**(Figure 4I** and **Figure S4C)**. Together these data suggest that the circulating SASP milieu impairs NO production and promotes excess mitochondrial superoxide bioactivity in senescent endothelial cells and these SASP-mediated differences are driven by in part by CXCL12 and may be prevented by fisetin supplementation.

### Senolytic treatment mitigates endothelial-to-mesenchymal transition (EndoMT) with aging

CXCL12 plays a crucial role in promoting the EndoMT, a transdifferentiation process by which endothelial cells lose their characteristic traits, while acquiring the contractile features of mesenchymal cells^36–38^. Importantly, the EndoMT is highly implicated in vascular aging and previous studies report shifts in senescent endothelial cells towards more contractile properties and reduced NO production^36,37^. As such, we next sought to determine if the EndoMT was an underlying mechanism mediating SASP- and CXCL12 -induced dysfunction in senescent endothelial cells. To do so, we first investigated whether senescent endothelial cells exhibit characteristics of EndoMT, and then assessed whether fisetin supplementation could prevent EndoMT induced by the circulating SASP and CXCL12. We initially measured transcripts specific to endothelial cells and mesenchymal cells in the senescent endothelial subcluster 10 and found that the old vehicle group demonstrated reduced endothelial characteristics and higher mesenchymal markers compared to the others groups **(Figure 5A)**.

**Figure 5.**
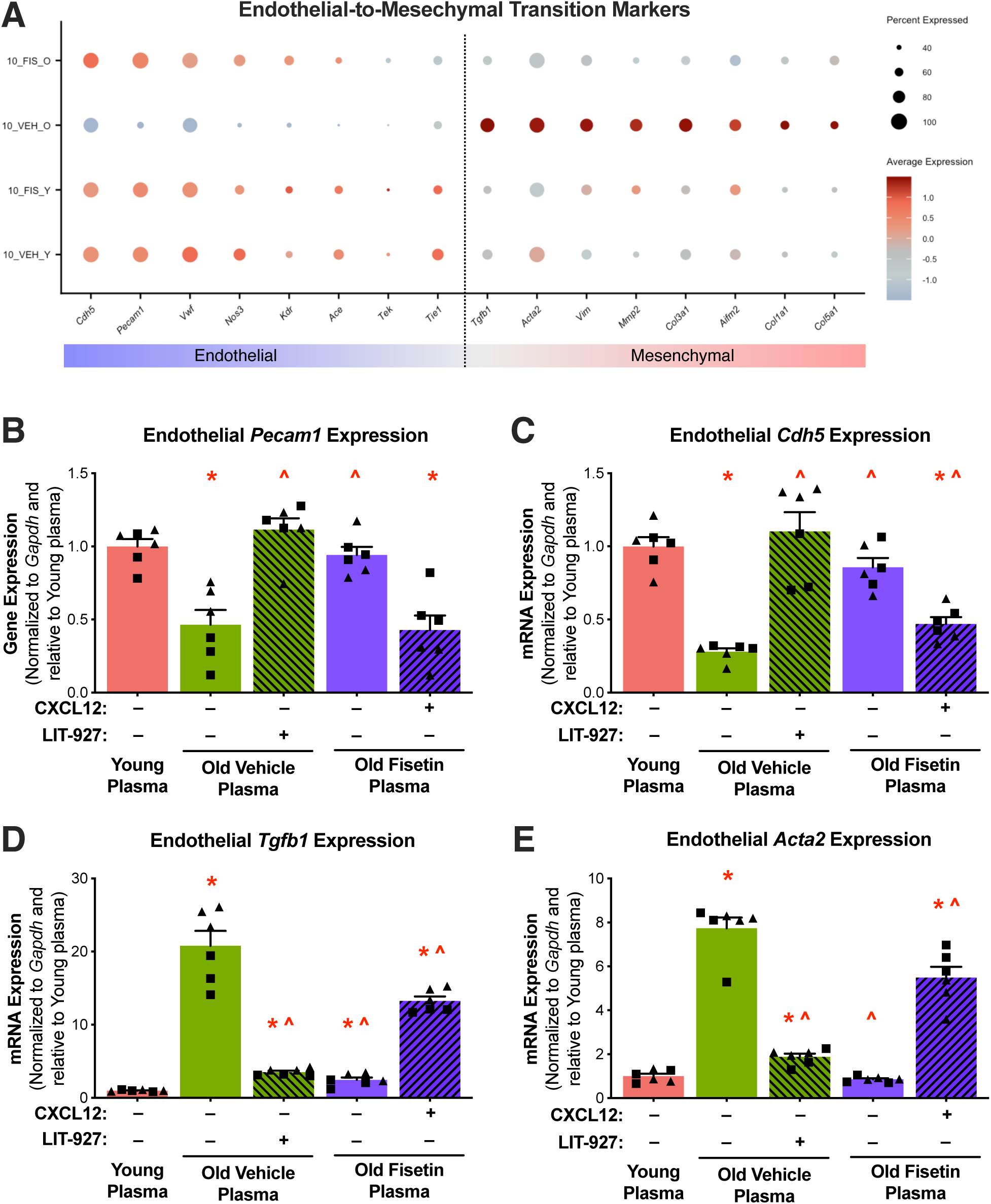
Fisetin supplementation prevents circulating SASP- and Cxcl12-induced endothelial-to-mesenchymal transition (EndoMT). Dot plot of endothelial and mesenchymal mRNA cell markers in senescent endothelial subcluster 10 **(A)**. mRNA levels of endothelial cell biomarkers *Cdh5* **(B)** and *Pecam1* **(C)** and mesenchymal cell biomarkers *Tgfb1* **(D)** and *Acta2* **(E)** in human aortic endothelial cells (HAECs) following plasma exposure with or without the addition of recombinant CXCL12 and its inhibitor, LIT-927. Values represent mean ± SEM; *p<0.05 vs. young control, ^p<0.05 vs. old vehicle; triangles represent females, squares represent males.

To determine whether the age-related circulating SASP milieu promotes EndoMT and whether these effects are mediated by CXCL12, we exposed HAECs to plasma from treated mice, either with CXCL12 added or inhibited. We then assessed the expression of the most affected endothelial markers (*Cdh5* and *Pecam1* mRNA) and mesenchymal markers (*Tgfb1* and *Acta2* mRNA). HAECs exposed to plasma from aged vehicle-treated mice showed significantly reduced mRNA expression of *Cdh5* (-72%, p<0.0001) and *Pecam1* (-54%, p<0.0001) compared to cells exposed to plasma from young mice. This age-related increase was mitigated by exposure to old fisetin plasma in *Cdh5* (3.1-fold, p<0.0001) and *Pecam1* (2-fold, p<0.0001) mRNA expression compared to old vehicle plasma **(Figure 5B** and **5C** and **Figure S5A** and **S5B)**. The addition of LIT-927 to old vehicle plasma increased *Cdh5* (4-fold, p<0.0001) and *Pecam1* (2.4-fold, p<0.0001) mRNA expression compared to old vehicle plasma alone **(Figure 5B** and **5C** and **Figure S5A** and **S5B)**. The addition of CXCL12 to old fisetin plasma reduced *Cdh5* (-45%, p<0.0001) and *Pecam1* (-55%, p<0.0001) mRNA expression compared to old fisetin plasma alone **(Figure 5B** and **5C** and **Figure S5A** and **S5B)**.

Contrary to the observed changes in endothelial markers, HAECs exposed to plasma from old vehicle mice had higher *Tgfb1* (21-fold, p<0.0001) and *Acta2* (7.7-fold, p<0.0001) mRNA expression compared to young plasma. Compared to old vehicle plasma, exposure to old fisetin plasma reduced *Tgfb1* (-88%, p<0.0001) and *Acta2* (-89%, p<0.0001) mRNA expression **(Figure 5D** and **5E** and **Figure S5C** and **S5D)**. The addition of LIT-927 to old vehicle plasma reduced *Tgfb1* (-83%, p<0.0001) and *Acta2* (-76%, p<0.0001) mRNA expression compared to old vehicle plasma alone **(Figure 5D** and **5E)**. The addition of CXCL12 to old fisetin plasma increased *Tgfb1* (5.4-fold, p<0.0001) and *Acta2* (5.4-fold, p<0.0001) mRNA expression compared to old fisetin plasma alone **(Figure 5D** and **5E** and **Figure S5C** and **S5D)**. In combination, these data indicate that the circulating SASP milieu and CXCL12 promote EndoMT in senescent endothelial cells and that these age-related changes in cell phenotypes are prevented by fisetin supplementation.

## DISCUSSION

In the present study, we assessed vascular cellular senescence at single-cell resolution in the context of aging and senolytic treatment with fisetin. Our findings demonstrate that endothelial cells are particularly susceptible to undergoing cellular senescence with aging and that fisetin, a naturally occurring senolytic compound, effectively eliminates senescent endothelial cells in old animals. In senescent endothelial cells and circulation, a collection of SASP factors increased with aging and was reduced by fisetin supplementation. Of these factors, *Cxcl12* mRNA emerged as the most highly differentiated with aging and fisetin supplementation. These changes in the circulating SASP milieu contributed to age-related endothelial dysfunction and the effects of the circulating SASP milieu were largely mediated by CXCL12. Fisetin-induced reductions in circulating CXCL12 contribute to the beneficial effects of fisetin on endothelial function by mitigating mitochondrial oxidative stress, improving NO production, and favorably modulating endothelial transdifferentiation. These results provided novel insights into the role of senescent cells and the SASP in vascular aging and the potential of senolytic therapy to mitigate age-related endothelial dysfunction.

Vascular endothelial dysfunction is a key risk factor, indicator, and predictor for CVD. Cellular senescence has emerged as a potent underlying mechanism of age-related endothelial dysfunction and a likely contributor to CVDs, including atherosclerosis^39^, hypertension^40^, and occlusive stroke^41^. Previous studies have characterized age-related increases in cellular senescence burden in aortic samples from mice^42^, primates^43^, and humans^44^. Excess senescent cells accumulate in the vasculature among various mammals to promote chronic, low-grade inflammation and excess oxidative stress through the SASP^45,46^. But despite this initial evidence of vascular health improvements with senolytic treatment, there is a lack of information on endothelial cell senescence *in vivo* and the effects of senolytic treatment on endothelial cells. In the present study, aortic senescent cell characterization identified endothelial cells to develop a high cellular senescence burden with aging, which was reduced by senolytic treatment with fisetin. Previous studies have demonstrated that fisetin reduces whole-aorta cell senescence burden *in vivo*^14^ as well as in cultured senescent endothelial cells *in vitro*^14,16^. At a single-cell level, our study explored the effects of cellular senescence burden with aging and following *in vivo* fisetin supplementation across multiple cell types and identified that endothelial cells undergo cellular senescence with aging and can be eliminated by *in vivo* senolytic treatment with fisetin. Importantly, the senescent endothelial cell population that we identified was involved in pathways related to CV pathologies and revealed a high expression of SASP-related transcripts. As such, our findings align with previous studies reporting the efficacy of senolytic agents in reducing senescent cell burden in aging tissues. However, this study provides the first evidence of the *in vivo* effects of senolytic treatment on vascular cell senescence at a single-cell level to reveal direct effects of aging and fisetin on senescent endothelial cells.

The circulating SASP milieu is composed of a heterogenous collection of pro-oxidant and pro-inflammatory factors that varies depending on cell origin and celluar senescence induction. As such, the circulating SASP milieu is considered a major mediator of the adverse effects of senescent cells, but direct evidence for the impact of the SASP on endothelial function remains minimal. Emerging evidence indicates the circulating SASP milieu as a driver of age-related endothelial dysfunction^10^; however, the specific component(s) of the circulation responsible for transducing impairments in endothelial function remains to be elucidated. One of the key findings in the present study is the elucidation of the role of the circulating SASP milieu as an underlying mechanism of age-related endothelial dysfunction. Our plasma exposure experiments in isolated arteries establish a causal role for the aged circulating SASP milieu in impairing endothelial function, with these age-related impairments being largely prevented in old animals that received senolytic treatment with fisetin.

We next sought to determine candidate factors in the circulating SASP milieu that may be responsible for mediating endothelial dysfunction with aging, as identification of these molecules would provide insight into mechanisms by which cellular senescence drives endothelial dysfunction and establish novel therapeutic targets. The chemokine encoded by *Cxcl12* mRNA emerged as a factor most highly upregulated in senescent endothelial cells with aging, with its expression reduced following fisetin senolytic treatment as determined by scRNA-seq. Additionally, the concentration of CXCL12 was elevated in the plasma of old animals and restored to young levels by fisetin supplementation. The addition and inhibition of CXCL12 in our plasma exposure experiments supports a causal role for CXCL12 in age-related endothelial dysfunction, as inhibition of CXCL12 in old vehicle plasma improved endothelial responses and adding exogenous CXCL12 to young plasma induced endothelial dysfunction. Moreover, the addition of CXCL12 to old fisetin plasma recapitulated age-related endothelial dysfunction, indicating that senolytic treatment with fisetin mitigated circulating SASP milieu-related endothelial dysfunction, in part, by targeting CXCL12. Collectively, these findings establish an altered circulating SASP milieu as an underlying mechanism by which senolytic treatment with fisetin improves endothelial function with aging.

The circulating SASP milieu stimulates endothelial dysfunction through underlying mechanisms that promote chronic inflammation and oxidative stress^9,38^. Notably, the pro-inflammatory circulating SASP milieu may promote cellular senescence systemically and in nearby cells, further promoting tissue-level dysfunction^8^. Senescent endothelial cells also contain dysfunctional mitochondria which are key regulators of the SASP^47^. Dysfunctional mitochondria in senescent endothelial cells also produce excess ROS, which scavenges NO to reduce its bioavailability^34^. Here, we showed that the exposure of young isolated mouse arteries and cultured arterial endothelial cells to the aged circulating SASP milieu induced humoral cellular senescence, reduced NO production, and stimulated mitochondrial oxidative stress. Importantly, these age-related manifestations were prevented in arteries and endothelial cells exposed to plasma from old animals supplemented with fisetin. These results suggest that fisetin favorably modulates the circulating SASP milieu to prevent vascular aging mechanisms that drive endothelial dysfunction.

Given that CXCL12 has been implicated in inflammatory^30,48^ and cellular senescence-related pathways^24,25^, its effects as a circulating SASP factor provide mechanistic insight into how endothelial dysfunction develops with aging. CXCL12 regulates the transduction of downstream signaling pathways involved in the EndoMT, a process whereby endothelial cells lose their inherent endothelial cell phenotypes and gain enhanced contractile properties ^29,30,48^. The circulating SASP milieu-mediated endothelial dysfunction was largely stimulated by independent effects of CXCL12 via humoral cellular senescence induction, reduced NO production, and stimulated mitochondrial oxidative stress. Previous studies demonstrate that endothelial cell-derived CXCL12 promotes atherosclerosis progression^30^ and our findings support that CXCL12 induces endothelial dysfunction likely through the EndoMT, a manifestation that is prevented by senolytic treatment. The ability of fisetin to ameliorate CXCL12-driven endothelial dysfunction further underscores its therapeutic potential in mitigating vascular aging.

## CONCLUSIONS

In conclusion, this study identifies that senescent endothelial cells accumulate with advancing age and promote the secretion of SASP factors into circulation. Senolytic intervention with fisetin effectively eliminated senescent endothelial cells and highlighted CXCL12 as a critical SASP factor that is increased with advanced age and reduced by senolytic treatment. Mechanistically, we uncovered that the circulating SASP milieu and CXCL12 induce cellular senescence, reduce NO production, promote mitochondrial oxidative stress, and stimulate EndoMT to confer endothelial dysfunction. By demonstrating that senolytic treatment with fisetin effectively reduces CXCL12 levels and improves endothelial function, our findings provide mechanistic insight into the therapeutic potential of senolytic compounds in vascular aging. Importantly, these findings may have important implications for ongoing clinical trials investigating the efficacy of fisetin supplementation for improving endothelial function in healthy mid-life and older adults (NCT06133634). These results provide physiological and mechanistic insight that support the development, establishment, and translation of senolytic therapies to improve cardiovascular health with aging.

## DATA AVAILABILITY

RNA sequencing data are deposited in GSE239591 (token:sfqjscwkrrezpyh) and GSE239602 (token: itypuqwkhhwtlit).

## AUTHOR CONTRUBUTIONS

SAM, KM, DT, MG, DRS, MJR, ABH, and ZSC designed the study. SAM, KM, DT, NSV, CH, ANO, RM, SD, KOM, SD, and ZSC performed the experiments. SAM, KM, DT, RM, SD, ABH and ZSC analyzed the data. SAM, KM, DT, SD, KOM, MG, DRS, MJR, ABH and ZSC interpreted the data and wrote the manuscript. All authors read and approved the final version of the manuscript.

## SUPPLEMENTAL METHODS

### Animals and experimental design

For the intervention period, young (6 months) and old (27 months) wildtype mice were randomly assigned to receive vehicle (10% ethanol, 30% PEG400 and 60% Phosal 50 PG) or fisetin (100 mg/kg/day in vehicle). For the young mice, 12 mice (6M/6F) received the vehicle and 12 mice (6M/6F) received fisetin, and for old mice, 8 mice (3M/5F) received vehicle and 10 mice (4M/6F) received fisetin. Treatment was administered via oral gavage using an intermittent dosing paradigm – one week on; two weeks off; one week on, as previously established senolytic dosing paradigm^1,2^. Mice were sacrificed one to two weeks following the final dose to rule out any acute effects of the compound as the terminal half-life of fisetin is ∼3.1 hours in plasma^3^.

### Aortic single-cell RNA sequencing

Aortic single-cell transcriptomic analysis was performed, as previously described^4^. Briefly, mice were anesthetized with ketamine and xylazine and blood was extracted using cardiac puncture. The whole aorta was collected after left ventricular perfusion with 10 ml of PBS and quickly transferred to cold PBS. To prepare a single-cell suspension, perivascular adipose tissue was removed and whole aortas from three mice (per sex and treatment group) were cut into ∼ 1-mm pieces and digested with an enzyme solution consisting of 10 mg/ml Collagenase II (Gibco) and 1 mg/ml elastase (Worthington Biochemical Corp.) in Dulbecco’s Modified Eagle Medium (DMEM; 3 ml/sample) for 15 min at 37°C in a tissue culture incubator. The cell suspension was strained through a 40-µm filter and centrifuged at 500 x g for 5 min. Cells were resuspended in DMEM supplemented with 15% fetal bovine serum (FBS) and 5 mM of EDTA and then stained with 2 µg/ml of propidium iodide for 5 min to assess cell viability and single, viable cells were sorted using Fluorescence-Activated Cell Sorting (FACS) Aria FUSION (BD Bioscience) for 10x Genomics library preparation. The single-cell libraries were prepared with Chromium Next GEM Single Cell 3’ Kit v3.1 (10x Genomics) with Chromium Next GEM Chip G Single Cell Kit (10x Genomics) according to manufacturer’s protocol with Chromium Controller (10x Genomics). Briefly, ∼10,000 single cells were used for GEM generation, and the cDNAs were then checked on the Agilent Bioanalyzer with High Sensitivity DNA kit (Agilent). Prepared cDNAs were then used for library preparation that were checked on the Agilent Bioanalyzer with DNA 1000 kit (Agilent Technologies). The libraries were sequenced with Illumina NovaSeq 6000 sequencer at a depth of 70,000-120,000 reads per cell. Sequencing data are deposited in the NCBI’s Gene Expression Omnibus repository GSE296698 (token: yrajmsyujxgjbql).

Single-cell RNA sequencing data were processed using Cell Ranger (version 8.0.1) with the mouse reference reference GRCm39-2024-A (10x Genomics). The obtained read count matrices were subsequently analyzed in R using the Seurat package, version 5.2.0 with default parameters in all functions, unless otherwise specified^5^. Standard quality control filtering was applied to each sample to eliminate low-quality cells and potential doublets from downstream analysis. Filtering also removed cells containing more than 25% mitochondrial RNAs, expressing fewer than 200 or greater than 7,000 transcripts (**Figure S1A** and **S1B**). RNAs that were detected in less than 3 cells were excluded from the analysis. For each sample, we analyzed RNA data with “LogNormalize” method followed by running the FindVariableFeatures function to select 2,000 most variable RNAs for dimensionality reduction. The FindIntegrationAnchors function was applied to choose anchors for data integration. After performing Principal Component Analysis (PCA), the top 40 PCA dimensions were determined by the ElbowPlot method and used to create the Uniform Manifold Approximation and Projection (UMAP) with the resolution parameter set to 0.3. Unsupervised clustering of the merged revealed 29 clusters (**Figure S1C**). Differentially expressed marker RNAs for each cluster were identified with the FindAllMarkers function, and the FindMarkers function was used to find differentially expressed genes between experimental conditions. The main cell types were identified using marker genes of each cluster in combination with data from literature (**Figure S1D** and **S1E**) and revealed 11 distinct cell types in the aorta (**Figure S1F-H**). To identify subclusters within the cell types, the analysis was rerun separately on the cells of each cluster and the UMAPs were created using the resolution 0.3. Senescent cells assessment scoring was performed with Seurat function AddModuleScore. Differentially abundant RNA (DAR) testing was performed with the following cutoffs: adjusted p-value < 0.05, average log2 fold change > 0.25, and minimum percentage of cells in either group > 0.1. Pathway analysis was performed using KEGG (version 111.0)^6^ and Reactome (version 90)^7^ and was based on DAR testing.

### Plasma protein quantification

For multiplex analysis, plasma was thawed and centrifuged at 16,000 x g for 4 min. Custom murine Luminex Assay kits were designed by R&D Biosystems to include the following analytes: CCL4, CCL7, CCL8, CXCL12, FGF Basic, GDF15, IGFBP3, PAI-1, S100A9, and TIMP4. Plasma was diluted 1:10 using the Calibrator Diluent RD6-52 provided in the kit. Standards (provided with the kit), blanks, and plasma were incubated with the microparticle cocktail for 2 h at 25°C, followed by incubation with Biotin-Antibody cocktail for 1 h. The final incubation lasted 30 min with Streptavidin-PE and shaking at 25°C prior to running the plate on the Bio-Rad Bioplex-200 Instrument. Each incubation was followed by washing 3 times with Wash Buffer (provided in the kit). Instrument settings were adjusted to the following: 50 µl sample volume, Bio-Plex MagPlex Beads (Magnetic), Double Discriminator Gates set at 8,000 and 23,000, low RP1 target value for the CAL2 setting, 50 count/region. The results were analyzed with the Bio-Plex Manager software.

### Plasma-mediated vascular endothelial function

To assess the role of the circulating SASP milieu (plasma) and CXCL12 on endothelial function, an *ex vivo* isolated artery model was leveraged, as previously described^8^. In brief, carotid arteries were excised from young (3-6 month), intervention-naïve wildtype mice and cannulated onto pressure myographs. Plasma collected from young vehicle, old vehicle and old-fisetin treated mice was diluted in a solution containing 5% sex-matched plasma, 1% penicillin-streptomycin antibiotic cocktail, and 94% modified Krebs buffer corrected to pH 7.3. The diluted plasma samples were perfused luminally through the pressurized arteries for 24 h prior to assessing endothelial function. Following plasma perfusion, endothelial function was measured endothelium-dependent dilation (EDD) and endothelium-independent dilation (EID) in response to increasing doses of acetylcholine (ACh) and sodium nitroprusside (SNP), respectively, as described previously^1,8^. In brief, after vessels were pre-constricted with phenylephrine (PE; 2 mM; Sigma-Aldrich), EDD was assessed by measuring increases in luminal diameter in response to increasing concentrations of ACh (1 × 10^-9^ to 1 × 10^-4^ M; Sigma-Aldrich). Following EDD, EID was assessed by measuring the increase in luminal diameter in response to increasing concentrations of SNP, an exogenous NO donor (1 × 10^-10^ to 1 × 10^-4^ M; Sigma-Aldrich). All dose-response data are presented as percent dilation relative to maximum diameter to account for differences in baseline vessel diameter.

### Arterial mitochondrial superoxide bioactivity

Arterial mitochondrial superoxide bioactivity was assessed using the mitochondrial-specific superoxide spin probe 1-hydroxy-4-[2-triphenylphosphonio-acetamido]-2,2,6,6-tetramethylpiperidine (mitoTEMPO-H; Enzo Life Sciences) by electron paramagnetic resonance (EPR) spectrometry, as previously described^1,9^. In short, 1 mm aortic rings were incubated in 5% sex-matched plasma from young, old vehicle- or old fisetin-treated mice in DMEM for 24h. Following plasma exposure, aortic rings were incubated in Krebs/HEPES buffer containing 0.5 mM mitoTEMPO-H at 37°C for 1 h. Samples were analyzed using aMS300 Xband EPR spectrometer (Magnettech).

### Cultured endothelial cell nitric oxide (NO) and mitochondrial superoxide bioactivity

Human aortic endothelial cells (HAECs; PromoCell; used at passage 3-4; female, age: 80 years, non-smoker, free from known CVD) were cultured in a 96-well glass bottom plate (CellVis) under standard culture conditions (37.5°C, 100% relative humidity, 5% CO2). HAECs were grown in basal media (Endothelial Cell Growth Medium-2 [EGM-2] BulletKit; PromoCell) supplemented with 5% plasma collected from young, old vehicle, and old fisetin-treated mice for 2 h. HAECs grown in basal media and 5% FBS were used as a control condition. HAECs were stained with the fluorescent probe Hoechst (nuclei stain; Thermo Fisher) and either 10 µM diaminorhodamine-4M AM (DAR-4M AM; Sigma-Aldrich; to quantify NO production) for 45 min or 5 µM MitoSOX (Thermo Fisher; to quantify mitochondrial superoxide bioactivity) for 30 min. Live HAECs were imaged at 20x using wide-field fluorescence microscopy (EVOS M7000 Imaging System; Thermo Fisher) under standard incubation conditions. HAECs stained with DAR-4M AM were imaged before and 6 min after the addition of 100 µM acetycholine (Sigma) to stimulate NO production^10^ and mitochondrial superoxide bioactivity was assessed under basal (unstimulated) conditions^11^. Images were quantified using Celleste 5.0 Image Analysis Software (Thermo Fisher) as previously described^10,11^ and normalized to the FBS condition on each plate to control for potential variation between plates.

### Senescence-associated β-galactosidase (SA-β-Gal) staining

Arterial and cultured endothelial cell SA-β-Gal staining was performed using the Senescence Detection Kit (Abcam) according to the manufacturer’s instructions. Briefly, 1 mm aortic rings or HAECs (described above) were incubated in 5% sex-matched plasma from young, old vehicle- or old fisetin-treated mice in DMEM or EGM-2, respectively, for 24 h. Following plasma exposure, aortas and HAECs were washed with PBS, fixed (fixing solution provided in the kit), and incubated with the X-gal solution overnight. Aortic rings were then washed, frozen in OCT compound, and stored at -80°C until the time of sectioning. Aortic samples were later sectioned (7 µm; Leica CM300, Leica Biosystems) and plated in poly-L-lysine coated slides. Aortas and HAECs were stained with DAPI overnight and images were captured using bright-field and fluorescent microscopy (EVOS M7000 Imaging System; Thermo Fisher) at 10X magnification and quantified using ImageJ, as described^12^.

### Cultured endothelial cell gene expression

mRNA gene expression was measured in HAECs following incubation with 5% plasma from young, old vehicle or old fisetin-treated mice cultured in EGM-2 for 24 h. Briefly, RNA was extracted using the RNeasy mini kit (Qiagen). cDNA was synthesized using the iScript cDNA synthesis kit (Bio-Rad Laboratories). Transcripts of *Cdkn2a, Cdkn1a, Cdh5, Pecam1, Tgfb1,* and *Acta2* were analyzed using a StepOnePlus Real-Time PCR System (Applied Biosystems) in 96-well plates and the Taqman OpenArray (Applied Biosystems) was used as a master mix, as described. SimpleSeq DNA sequencing (Quintara Biosciences) was used to validate PCR products.

### CXCL12 protein addition and inhibition

Recombinant mouse CXCL12 protein (R&D Systems, Cat. #460-SD-050/CF) in sterile PBS was added back to plasma from young and old fisetin-treated mice, such that the concentration of CXCL12 matched the average levels in old vehicle-treated mice (3210 pg/ml) for CXCL12 add-back experiments. 1 µg/ml of LIT-927 (Selleckchem, Cat. #S8813)^13^ in DMSO was added to the old vehicle plasma for CXCL12 inhibition experiments.

## SUPPLEMENTAL FIGURES

**Figure S1.**
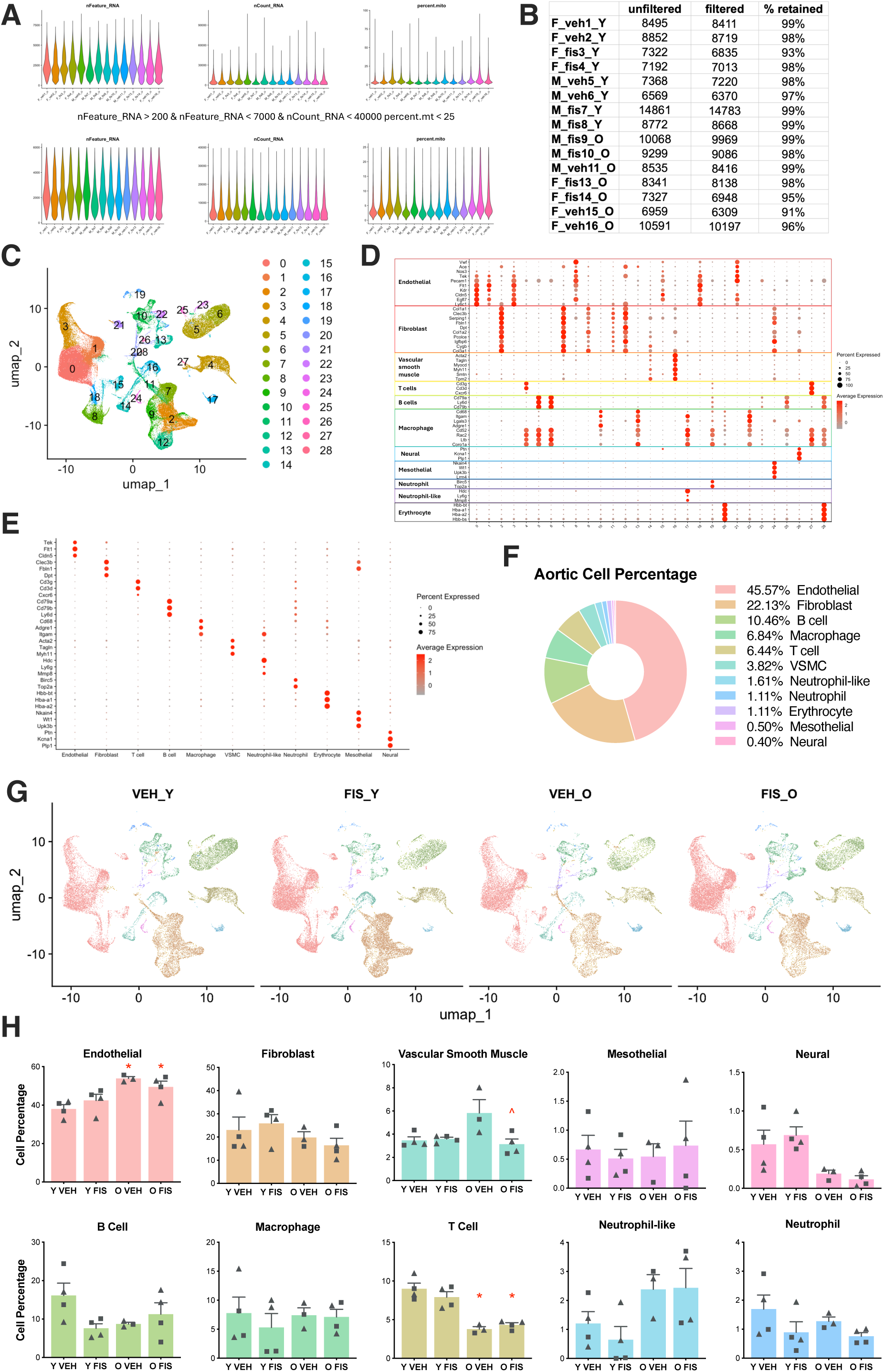
Whole aortic single-cell RNA sequencing quality control. Standard quality control filtering was applied to each sample to remove cells containing more than 25% mitochondrial RNAs, expressing fewer than 200 or greater than 7,000 transcripts **(A-B)**. Uniform Manifold Approximation and Projection (UMAP) of merged samples **(C)**. Cell types were identified using marker genes of each cluster **(D)**. Cell types were merged and validated by top 3 gene markers **(E)**. Aortic cell composition on merged samples by cell type **(F)**. UMAP **(G)** and cell composition analysis **(H)** of samples by cell type and treatment conditions: young vehicle (Y VEH), young fisetin (Y FIS), old vehicle (O VEH), and old fisetin (O FIS). Values represent mean ± SEM; *p<0.05 vs. Y VEH, ^p<0.05 vs. O VEH. Triangles represent females, squares represent males.

**Figure S2.**
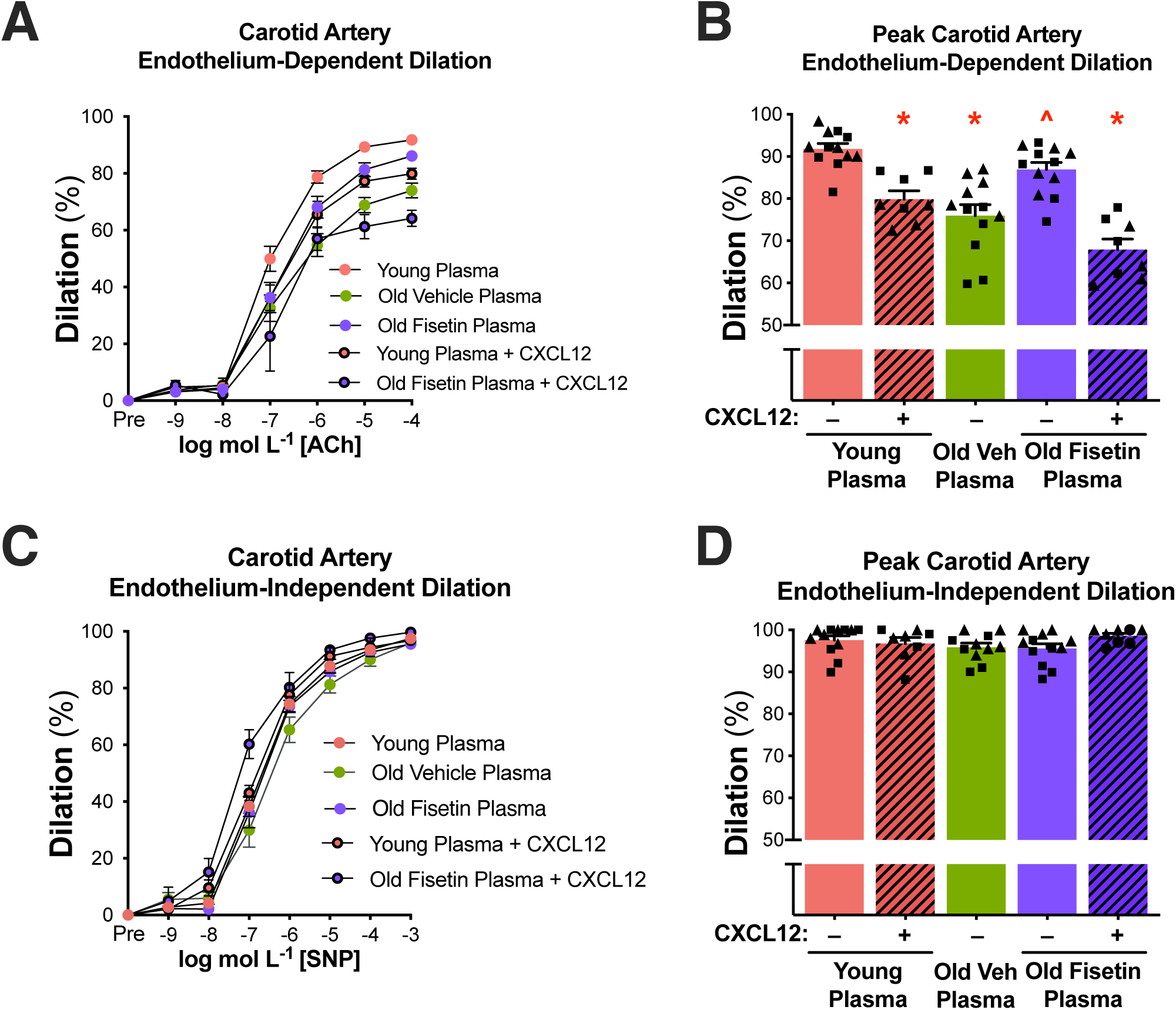
The circulating SASP- and CXCL12-induced endothelial dysfunction with aging and prevention by senolytic treatment with fisetin and CXCL12 inhibition. Endothelium-dependent dilation curves **(A)** and peak values **(B)** and endothelium-independent dilation curves **(c)** and peak values **(d)** in isolated carotid arteries from young mice following plasma perfusion with and without the addition of recombinant mouse CXCL12. Values represent mean ± SEM; *p<0.05 vs. young control, ^p<0.05 vs. old vehicle; triangles represent females, squares represent males. ACh: acetylcholine, SNP: sodium nitroprusside.

**Figure S3.**
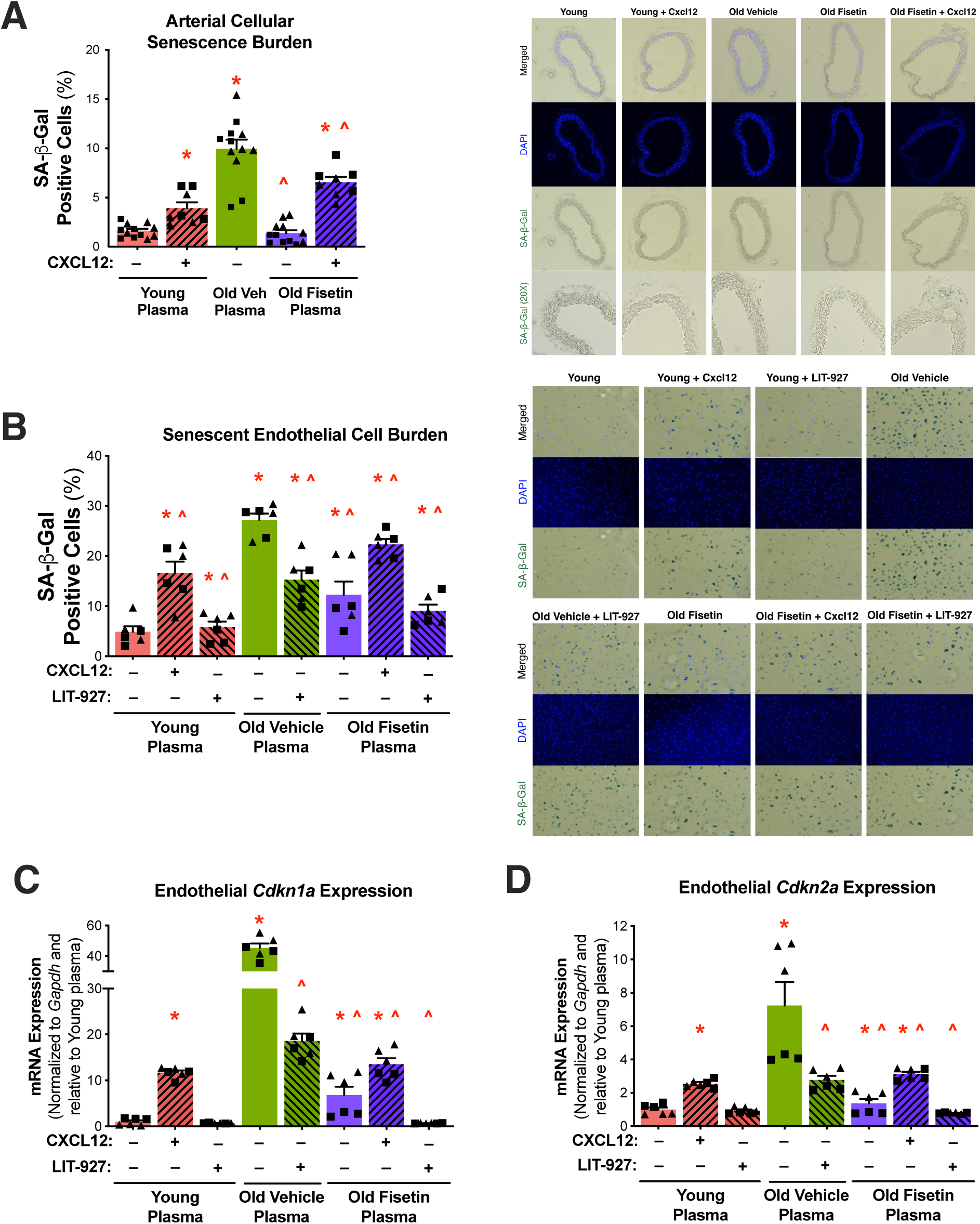
The circulating SASP- and CXCL12-induced cellular senescence with aging and prevention by senolytic treatment with fisetin and CXCL12 inhibition. Senescence-associated β-Galactosidase (SA-β-Gal) signal following plasma exposure with and without the addition of recombinant CXCL12 and CXCL12 inhibitor, LIT-927 in isolated arteries **(A)** and human aortic endothelial cells (HAECs) **(B)**, with representative images. Cellular senescence biomarkers *Cdkn1a* (C) and *Cdkn2a* (D) mRNA in HAECs following plasma exposure with and without the addition of recombinant CXCL12 and CXCL12 inhibitor, LIT-927. Values represent mean ± SEM; *p<0.05 vs. young control, ^p<0.05 vs. old vehicle; triangles represent females, squares represent males.

**Figure S4.**
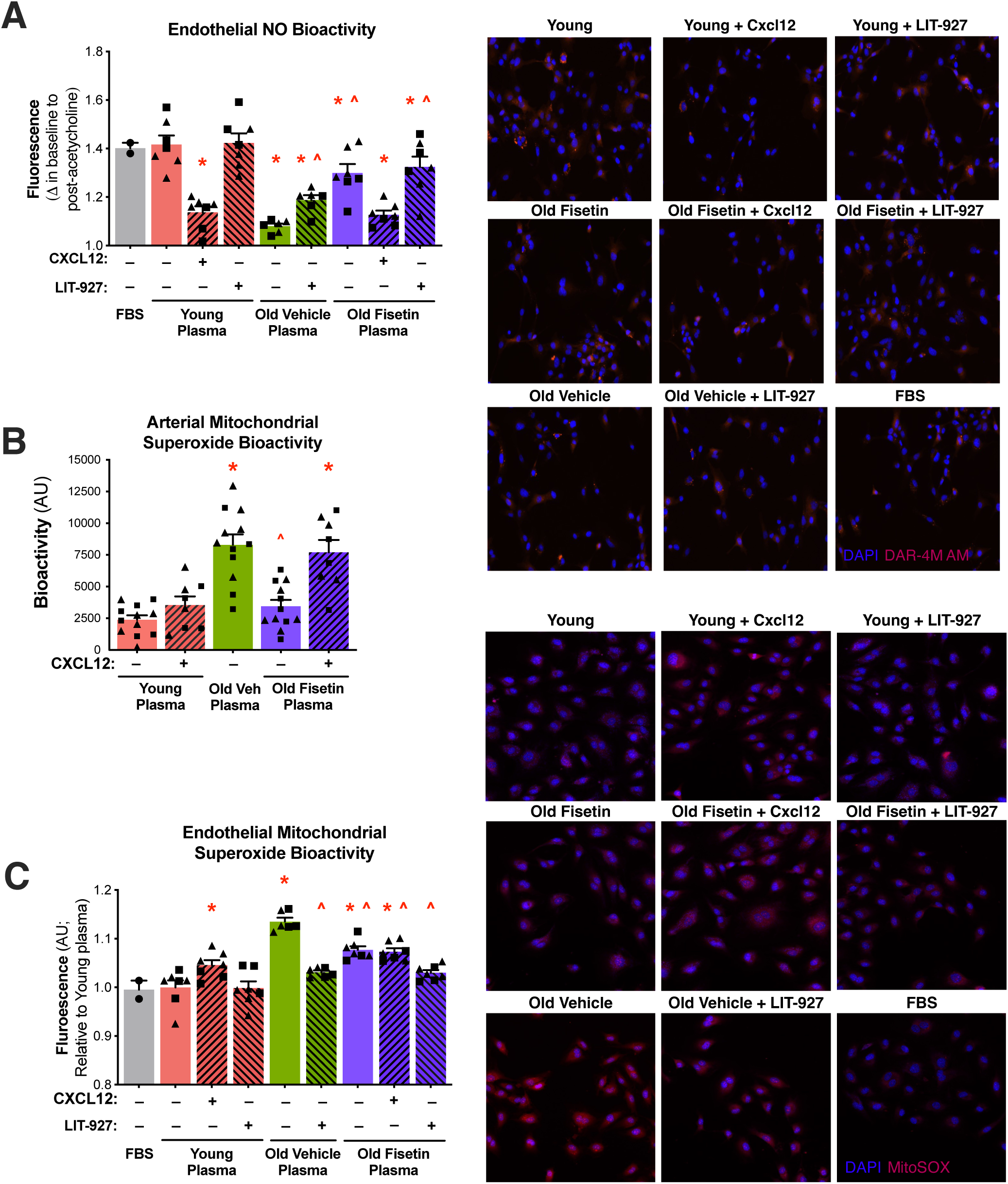
The circulating SASP- and CXCL12-induced acetylcholine-stimulated nitric oxide (NO) production and mitochondrial superoxide bioactivity with aging and prevention by senolytic treatment with fisetin and CXCL12 inhibition. NO bioactivity in human aortic endothelial cells (HAECs) following plasma exposure with and without the addition of recombinant CXCL12 and LIT-927 **(A)**. Mitochondrial superoxide bioactivity following plasma exposure with and without the addition of recombinant CXCL12 and LIT-927 in isolated arteries **(B)** and HAECs, with representative images **(C).** Values represent mean ± SEM; *p<0.05 vs. young control, ^p<0.05 vs. old vehicle; triangles represent females, squares represent males. FBS: fetal bovine serum, AU: arbitrary units.

**Figure S5.**
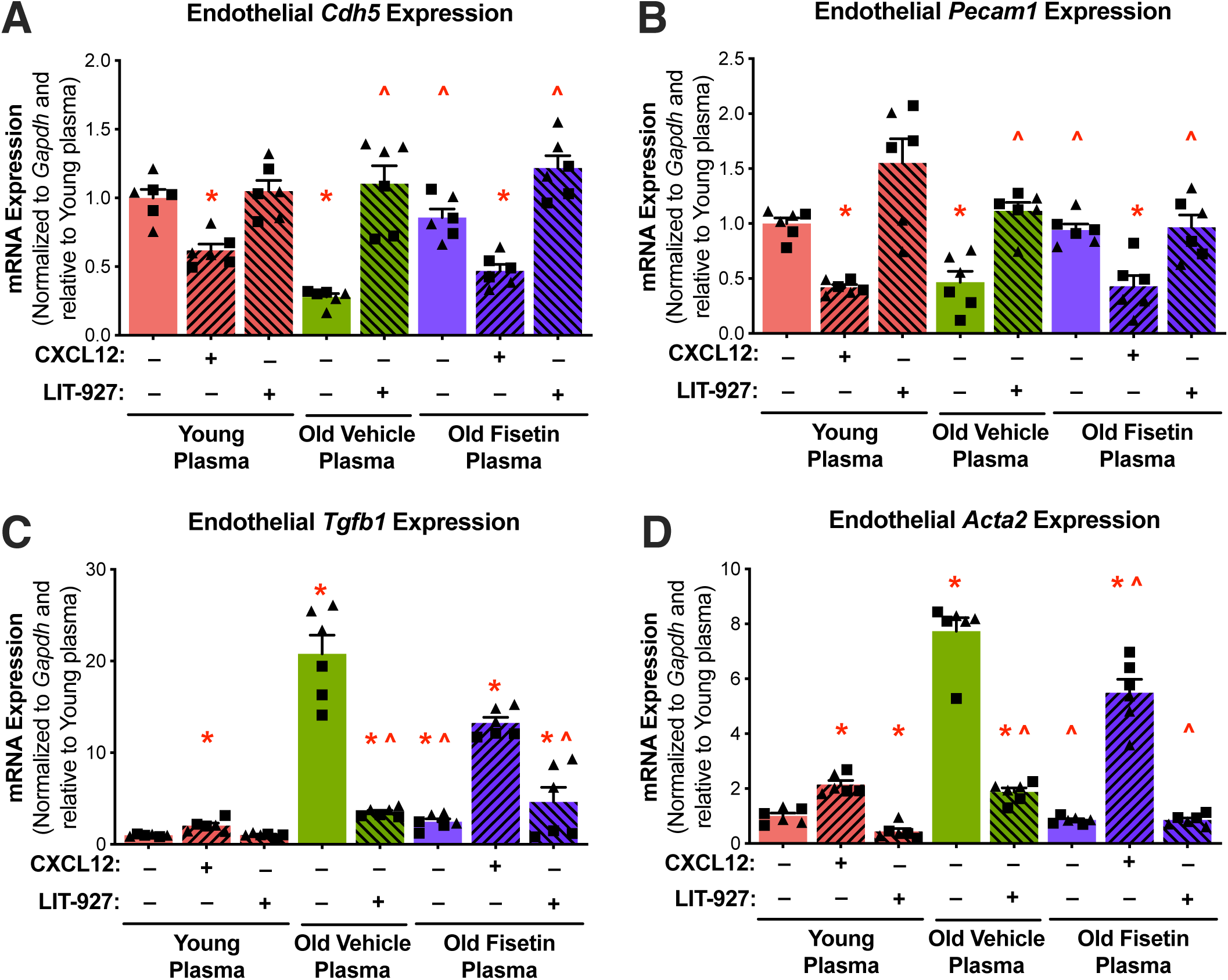
Fisetin supplementation prevents circulating SASP- and CXCL12-induced endothelial-to-mesenchymal transition. mRNA levels of endothelial cell biomarkers *Cdh5* **(A)** and *Pecam1* **(B)** and mesenchymal cell biomarkers *Tgfb1* **(C)** and *Acta2* **(D)** in human aortic endothelial cells (HAECs) following plasma exposure with or without the addition of recombinant CXCL12 and CXCL12 inhibitor, LIT-927. Values represent mean ± SEM; *p<0.05 vs. young control, ^p<0.05 vs. old vehicle; triangles represent females, squares represent males.

